# The CUT&RUN Greenlist: genomic regions of consistent noise are effective normalizing factors for quantitative epigenome mapping

**DOI:** 10.1101/2023.10.26.564165

**Authors:** Fabio N. de Mello, Ana C. Tahira, Maria Gabriela Berzoti-Coelho, Sergio Verjovski-Almeida

## Abstract

Cleavage Under Targets and Release Using Nuclease (CUT&RUN) is a recent development for epigenome mapping, but its unique methodology can hamper proper quantitative analyses. As traditional normalization approaches have been shown to be inaccurate, we sought to determine endogenous normalization factors based on regions of constant nonspecific signal. This constancy was determined by applying Shannon’s information entropy, and the set of normalizer regions, which we named the “greenlist,” was extensively validated using publicly available datasets. We demonstrate here that the greenlist normalization outperforms the current top standards, and remains consistent across different experimental set-ups, cell lines, and antibodies; the approach can even be applied to other organisms or to CUT&Tag. Requiring no additional experimental steps and no added cost, this approach can be universally applied to CUT&RUN experiments to greatly minimize the interference of technical variation over the biological epigenome changes of interest.

## Introduction

With the advent and popularization of high-throughput sequencing technologies, the field of genome biology experienced drastic changes as several genomic assays were adapted to high-throughput approaches. Chromatin immunoprecipitation and sequencing (ChIP-seq) is perhaps the most popular among them [1–3], allowing for target-specific isolation of DNA-protein complexes, and thus the genome-wide mapping of epigenetic modifications, chromatin-modifying enzymes, and transcription factors. Similarly, Cleavage Under Targets and Release Using Nuclease (CUT&RUN) is a recent development by Skene & Henikoff [4] for genome mapping of DNA-protein interactions, an optimization of chromatin immunocleavage (ChIC) [5] for high-throughput sequencing, which seeks to specifically address some of ChIP-seq’s main drawbacks. Rather than relying on the random shearing and immunoprecipitation of chromatin, CUT&RUN relies on a micrococcal nuclease (MNase) protein fusion guided by a protein-A/G-conjugated antibody, directing the cleavage activity to the genomic loci of interest. This approach greatly minimizes the generation of antibody-nonspecific fragments, i.e. noise [4, 6]. In turn, this significantly higher signal-to-noise ratio results in much lower requirements for both starting sample volumes and required read depths [4, 6, 7], greatly reducing the sequencing cost compared to ChIP-seq. Additionally, this target-directed enzymatic cleavage enhances the accuracy [4, 7], as it is no longer necessary to rely on random shearing from sonication or undirected enzymatic cleavage.

The advantages of CUT&RUN have led to its increasing adoption [8–10], gaining popularity as a simpler, cheaper alternative to ChIP-seq. A derived technique has been developed, Cleavage Under Targets and Tagmentation (CUT&Tag) [11], by integrating Tn5 transposase tagmentation, resulting in an even higher signal-to-noise ratio.

Early CUT&RUN mainly relied on the numerous tools and analysis concepts previously developed for ChIP-seq [4]. However, the methodological differences of CUT&RUN have since prompted the development of specialized tools and approaches, such as specialized peak calling algorithms [7, 12] and quality control analysis [13, 14].

A recurrent challenge has been proper signal normalization to make samples comparable; for ChIP-seq analyses, straightforward library size-based approaches (such as fragments per kilobase million or fraction reads in peaks) have been extensively found to be inappropriate [15–17]. Normalizing by nonspecific noise can be an option, however CUT&RUN’s low noise generation typically results in an inconsistent background [4, 7], inappropriate for this kind of approach. So far, the accepted golden standard has been spike-in normalization [4, 6, 16, 18]; however, this approach fails to account for the cleavage efficiency and patterns, as the spike material is separately fragmented in advance [17, 19, 20].

In the course of our work with CUT&RUN (unpublished) we began to search for better alternatives, building upon concepts presented in the ENCODE blacklist of problematic regions for ChIP-seq [21]. This work identified that certain regions of the genome present very high background signals, regardless of other experimental factors, being reliably used in ChIP-seq analyses [21–23].

In the present work, we aimed to expand upon this concept by identifying a set of genomic regions that exhibit consistent background signals in CUT&RUN experiments. We have coined the term “greenlist” to refer to this list of regions, following the naming scheme used for ChIP-seq blacklists [21], greylists [24], and sequencing barcoding whitelists. We demonstrate that these greenlist signals remain consistent across various CUT&RUN experiments involving different antibodies and cell types, thus making them suitable for use as control signals for normalization purposes. We have made available a human CUT&RUN greenlist and blacklist, a mouse CUT&RUN greenlist and blacklist, as well as a human CUT&Tag greenlist and blacklist as Supplementary Material S1. This provides a robust, intrinsic solution for quantitative CUT&RUN analysis, surpassing current methods without any additional cost or experimental steps.

## Results

### Generating the Greenlist

Similar to the ENCODE blacklist [21], we sought to develop a systematic pipeline to identify genomic regions with constant signals across all publicly available negative control CUT&RUN samples. After filtering, we obtained 463 human samples from 102 experiments, derived from 73 established cell lines and 112 patient biopsy samples, and 30 commercial anti-IgG antibodies (plus 56 samples with no antibody); this variety is essential in ensuring that our results are not biased to specific cell types or experimental setups. As expected, we observed variable correlations between samples (Fig. 1a), especially across different experiments. To better understand these differences, we performed a principal component analysis (Fig. 1b) to test the correlation of the components to known meta-factors, which revealed that this variation was mainly related to differences in protocol/commercial kit used, cell line, and sequencing depth (Fig. 1c, Supplementary Data S2). We also saw significant variation across experiments not attributed to other tested meta-factors (Fig. 1c), indicating that there are still experiment-specific sources of noise of unknown origin. In fact, we expect a fair portion of noise generation to be stochastic in nature, and not all variation to be fully explainable.

**Figure 1:**
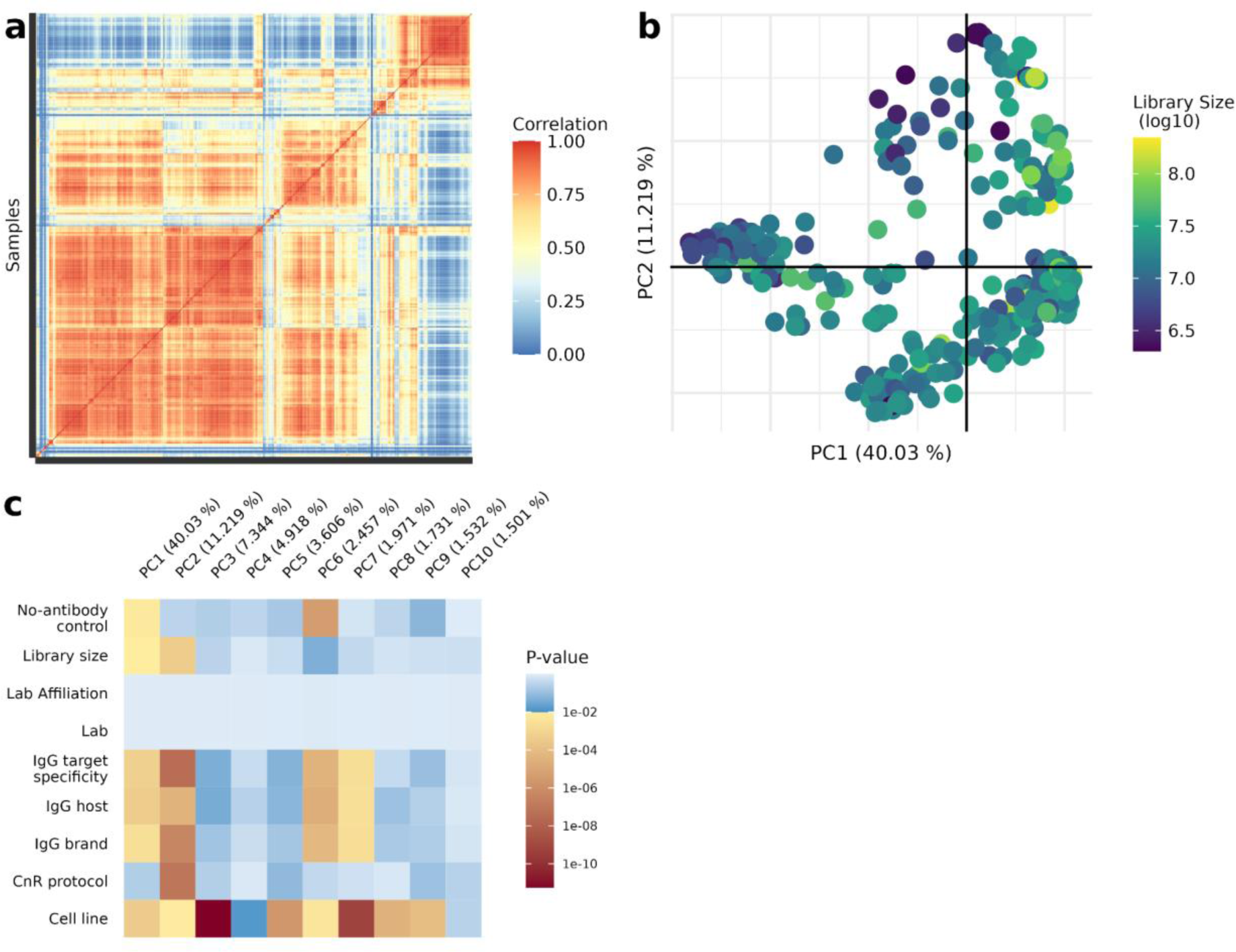
Overview of publicly available negative control CUT&RUN samples. Comparison of the 463 CUT&RUN samples selected for analysis. **a,** Heatmap of Pearson correlation between the samples, considering all genomic bins (1kb); **b,** First two components of the PCA, colored by aligned library size; **c,** p-value of association between each of the first 10 principal components (PC1 to 10, columns) to known meta-factors (lines), tested by ANCOVA; p-values considered significant (p≤0.01) are colored in shades of yellow to red.

Of note, the processing of these datasets confirmed the ubiquity of CUT&RUN’s varying yields, with several datasets presenting extreme variations in library size (up to 48x difference), even across supposedly identical replicates, as well as samples presenting very low alignment percentages (1.34-99.78%). These differences can have a major impact on quantitative analyses, highlighting the need for better quantitative tools.

To identify how constant the signal of each region is across all the samples, we relied on Shannon entropy [31,32] (Fig. 2a). Briefly, we expect bins with inconsistent high-count outliers to have low information entropy, and bins with a more homogenous distribution across the dataset to have higher entropy (further detailed in the Methods). This approach is preferable to simply using the standard deviation or standard error, as those would require the assumption that all bins follow the same general distribution function, which cannot be assumed in advance. As expected, progressively filtering out the lowest entropy bins greatly increased the correlation between samples (Fig. 2b), indicative of how consistent the signals from the high entropy regions were.

**Figure 2:**
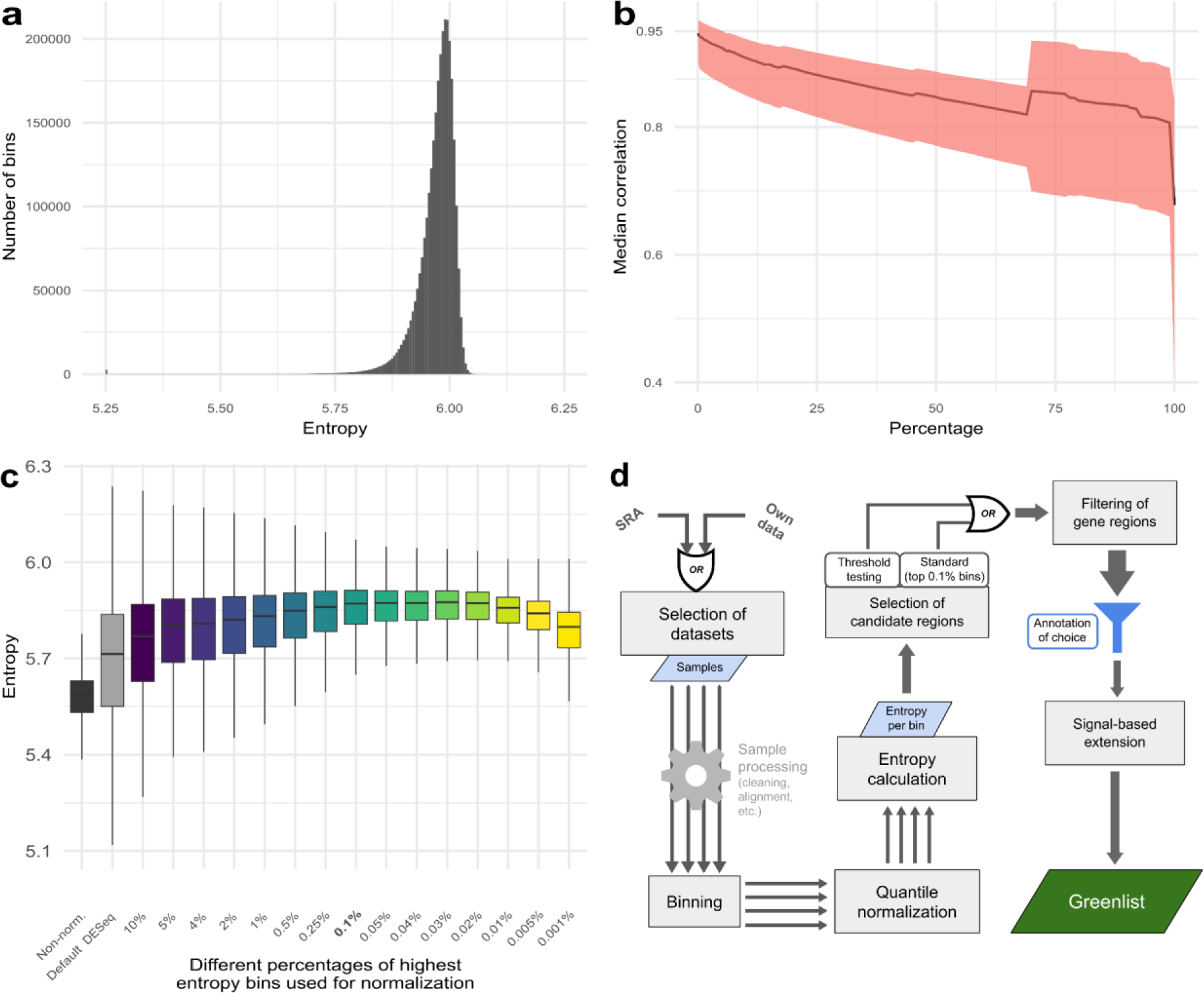
High entropy regions are effective normalizers. Shannon entropy calculations over all genomic bins (excluding those with no counts on any samples). **a,** Frequency distribution of entropy for all bins; **b,** Median Pearson correlation between samples (line), considering different fractions of the highest entropy bins, with Q25 to Q75 range shown in red; **c,** Entropy distributions for our test set (the 10% of bins with the lowest entropies) after normalization using different percentages of highest entropy bins, as indicated in the x-axis; in the y-axis, boxplot represents median entropy, first and third quartiles, whiskers extend to 5% and 95% quantiles; **d,** Summary of the processing pipeline used for greenlist generation, generalized for applicability in new contexts.

We computed the entropy distributions of our test set (equivalent to 10% of bins) after normalization using candidate regions with progressively smaller percentages of highest entropy bins and compared them with a standard DESeq normalization based on library size (Fig. 2c). We also compared with the entropy computed with non-normalized counts, which was very low (Fig. 2c), as technical variation and different sequencing depths compound with the biological variation and lead to extreme differences between the samples. A nearly ideal threshold was approached by using normalization regions with progressively smaller percentages of highest entropy bins (Fig. 2c); a perfect normalization would maximize entropy, nullifying technical variation and leaving only the biological. Based on these results, we selected the top 0.1% most entropic bins as the threshold for generating the greenlist (Fig. 2c, highlighted). This threshold maximized entropy and avoided overfitting, whereas the normalization with very small fractions of highly entropic bins (< 0.02%) decreased entropy, as more bias was introduced by enhancing the effect of minor stochastic variations, being less representative of the overall data.

We subsequently filtered out from the candidate greenlist any region overlapping or close to any known genes (< 5kb away from known genes), to avoid possible overlapping of true signals, i.e., fragments generated in an antibody-specific manner within gene bodies and neighborhoods. Finally, we extended and merged a selected region if other regions in its vicinity were in the top 1% highest entropy bins (similarly to the original blacklist [21]) in order to avoid short, scattered regions, thus obtaining our CUT&RUN greenlist (available as Supplementary Material S1). The pipeline for greenlist generation is summarized in Figure 2d.

Evaluating this final list by the same previous parameters, we observed results comparable to the initial threshold tests (Fig. 3a-b). Some loss of efficiency is to be expected, since many high-entropy regions are lost when filtering out gene neighborhoods, but this extra cautious approach did not seem to significantly affect the performance of the final list. To ensure this entropy maximization behavior wasn’t simply due to the size of our greenlist, we performed a Monte-Carlo simulation randomly selecting 0.1% of bins (Fig. 3c) and saw that our greenlist normalization showed a significantly (p<10^-6^, 100,000 trials) higher entropy median than a random selection, both before and after filtering out gene neighborhoods. We also confirmed that the signal from these regions did not seem to be significantly biased to experimental differences such as different antibodies or cell types (Fig 3d, Supplementary Material S2), with each principal component only explaining small fractions of the total variance (Fig. 3e-f). We did observe a significant association (p<10^-17^) between library size and the first principal component (even after normalization), but that was expected: higher sequencing depths will highlight subtle variations along the regions, while lower depths will mask them. Regardless, this accounted for only 4% of the total variation of the dataset (Fig. 3f). Some of the remaining principal components were affected by cell type, suggesting that greenlist regions can still be slightly impacted by differences in chromatin accessibility. Importantly, differences in experimental setups, such as different antibody brands/isotypes, protocol optimizations, or test variables of interest, showed limited impact on the overall observed variability (< 25% counting the 20 PCs) (Fig 3d,3f). These greenlist regions also showed no significant distribution bias along genomic features (Supplementary Materials S3).

**Figure 3:**
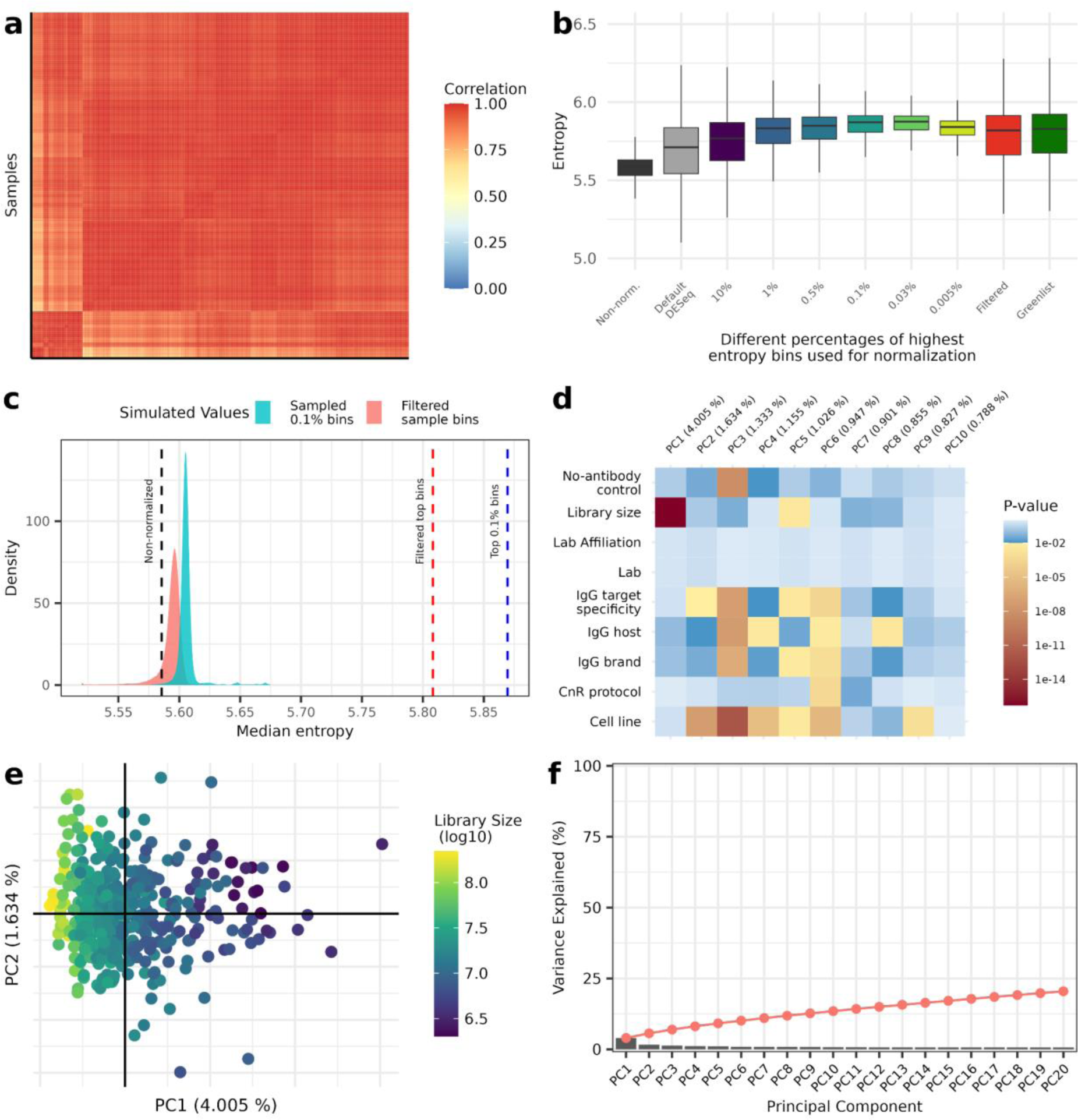
Greenlist regions show consistent signal across variate experimental conditions. Evaluation of the performance of the final greenlist for human CUT&RUN, after filtering and extension. **a,** Heatmap of Pearson correlation across all samples, considering only greenlist regions signal; **b,** Performance of final greenlist (dark green) and greenlist without extension (red) as normalizers of our test set (previous results similar to Figure 2c are shown); **c,** Density distribution of median entropies of Monte-Carlo simulations of normalizations with randomly selected regions, before (blue) and after (red) filtering (medians obtained in Figure 3b shown as dashed lines); **d,** p-value of association between each of the first 10 principal components (PC1 to 10, columns) to known meta-factors (lines), tested by ANCOVA (p-values considered significant (p≤0.01) are colored in shades of yellow to red); **e,** First two components of PCA of greenlist regions across all samples, color-coded according to library size, the only significant (p≤0.01) cofactor found; **f,** Scree plot of each PCA component’s contribution to total variance, cumulative variance shown as red line.

### A robust approach for different organisms, CUT&RUN or CUT&Tag

Next, we sought to expand the applicability of our greenlist by defining the greenlist for the mouse genome. Application of the pipeline to 611 mouse negative control samples showed similar quality metrics as seen before (Fig. 4a, Extended Data 4a-b), indicative of the consistency of the method. Next, we asked whether a viable greenlist could be generated from a single experiment with a large enough sample size; this should cover cases where a greenlist is needed for an organism with none or few previously available public samples. For this, we used the GSE151326 GEO dataset [33], an antibody characterization study with 50 samples of 43 different antibodies on human cells, as a demonstration. We see that despite how different the samples were initially, filtering the highest entropy regions yielded very consistent signals (Fig. 4b, Extended Data 4c), which did not appear to be biased by library size or antibody target. Comparing this *de novo* greenlist with our previous hg38 greenlist, we observed a middling correlation of the entropy metrics (Fig. 4c) but minimal overlap of the final lists (Fig. 4d), as measured by precision-recall F1 scores. This is a direct consequence of the variety of cell types used for the original, which are lacking in a single experiment-based list; nevertheless, the consistency of the found *de novo* greenlist regions shows that this would be a suitable normalization option for the experiment at hand, just not as widely applicable as one built from several experiments.

**Figure 4:**
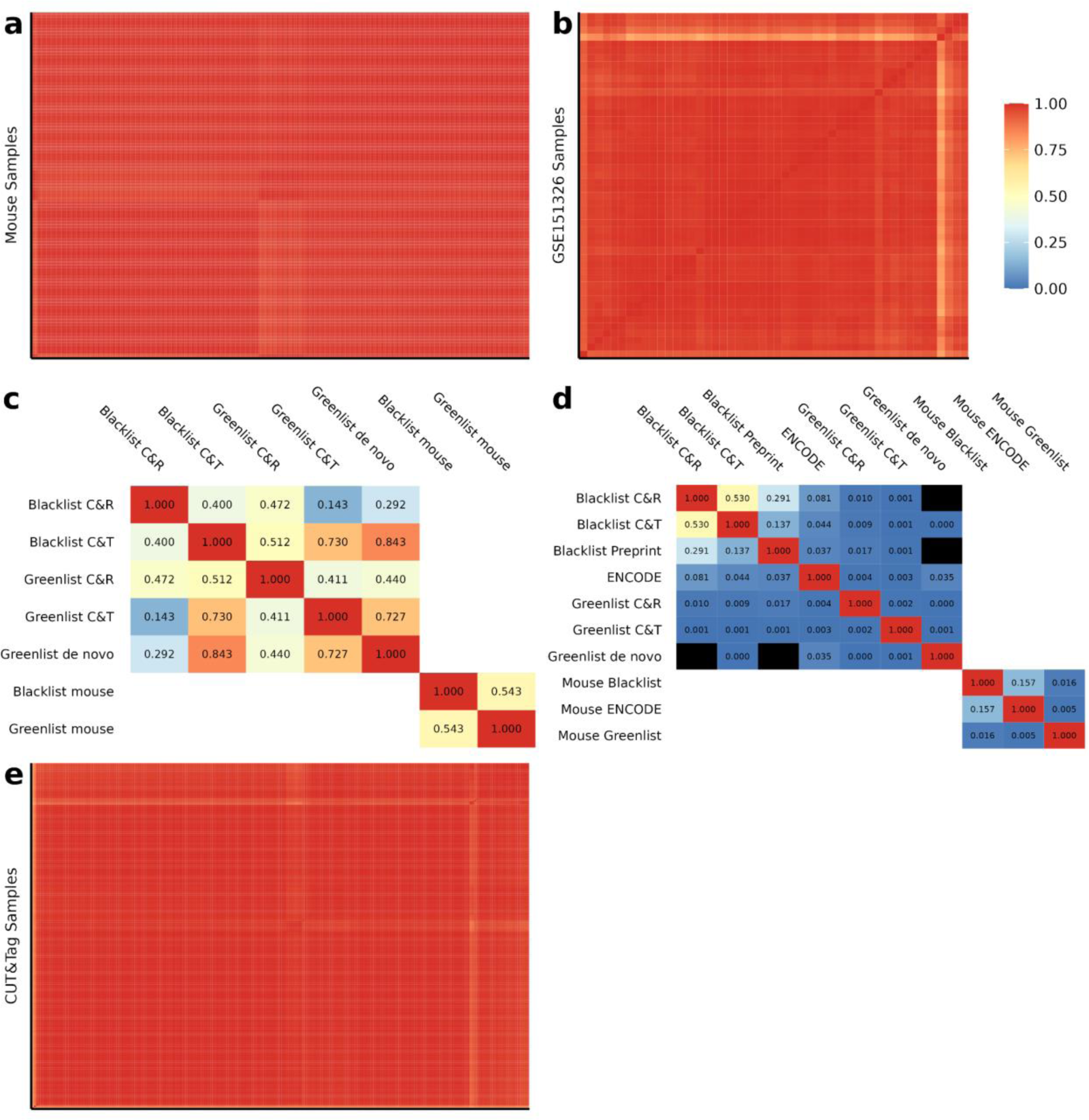
Other applications of our pipeline show similar performances. **a,** Heatmap of Pearson correlation of 611 mouse CUT&RUN samples, considering only signal from greenlist regions; **b,** Heatmap of Pearson correlation of the 50 samples from dataset GSE151326, considering only signal from the *de novo* greenlist constructed off this dataset alone; **c,** Spearman correlation between the quantifications used for creating each list, either entropy (for greenlists) or median signal (for blacklists), considering all genomic 1kb bins; **d,** Overlap between the final lists created and some from the literature, considering F1 scores of precision-recall; **e,** Heatmap of Pearson correlation of 217 human CUT&Tag samples, considering only signal from the CUT&Tag greenlist regions. All graphs share the same color scale (displayed near b).

Additionally, we also expanded our efforts towards CUT&Tag. Despite intrinsically producing even less background noise than CUT&RUN, the analysis of 217 human CUT&Tag negative control samples still revealed a greenlist of consistent signal regions (Fig. 4e, Extended Data 4d), featuring similar entropy maximization results as observed for CUT&RUN (Extended Data 4e). Comparing the two lists we again saw little overlap (Fig. 4d), suggesting that each list is accurately optimized to their respective technique.

In parallel, we also took the opportunity to create blacklists for human CUT&RUN, mouse CUT&RUN, and human CUT&Tag, following the methods of the original ChIP-seq blacklist [21]. Despite having since been validated for other techniques, the ENCODE ChIP-seq blacklist has not been validated for CUT&RUN or CUT&Tag, and the vastly different methodological steps (ie. the lack of random fragmentation and immunoprecipitation) may affect the generation of high-signal background regions. We observed a fair overlap between the CUT&RUN and CUT&Tag blacklists (Fig. 4d), indicative of the similarity of these techniques, but as expected both feature little overlap with the ChIP-seq blacklist, confirming their drastically different noise profile. Previous efforts have been made to create a CUT&RUN-specific blacklist [25] but with limited samples (N=20), which would present limited accuracy. Importantly, we observe minimal overlap between the greenlists and blacklists for each method.

### Greenlist normalization outperforms current standards

We have observed that the greenlist regions act as suitable normalizing factors in our tests, maximizing the entropy of our negative control datasets, so we next sought to compare them to consolidated normalization approaches. We reanalyzed dataset GSE104550 [26], generated by the Henikoff lab and used by Meers *et al.* [6] to establish the comparability of normalizations by *E. coli* carryover spike-in and external spike-in (such as added *S. cerevisiae* DNA), the two normalization methods considered as the golden standards for CUT&RUN. Here we again observe that the signal from greenlist regions remained consistent, despite the overall drastic differences between sample profiles (Fig. 5a-b), which are mainly due to the two antibodies’ (anti-CTCF and anti-H3K27me3) different binding profiles.

**Figure 5:**
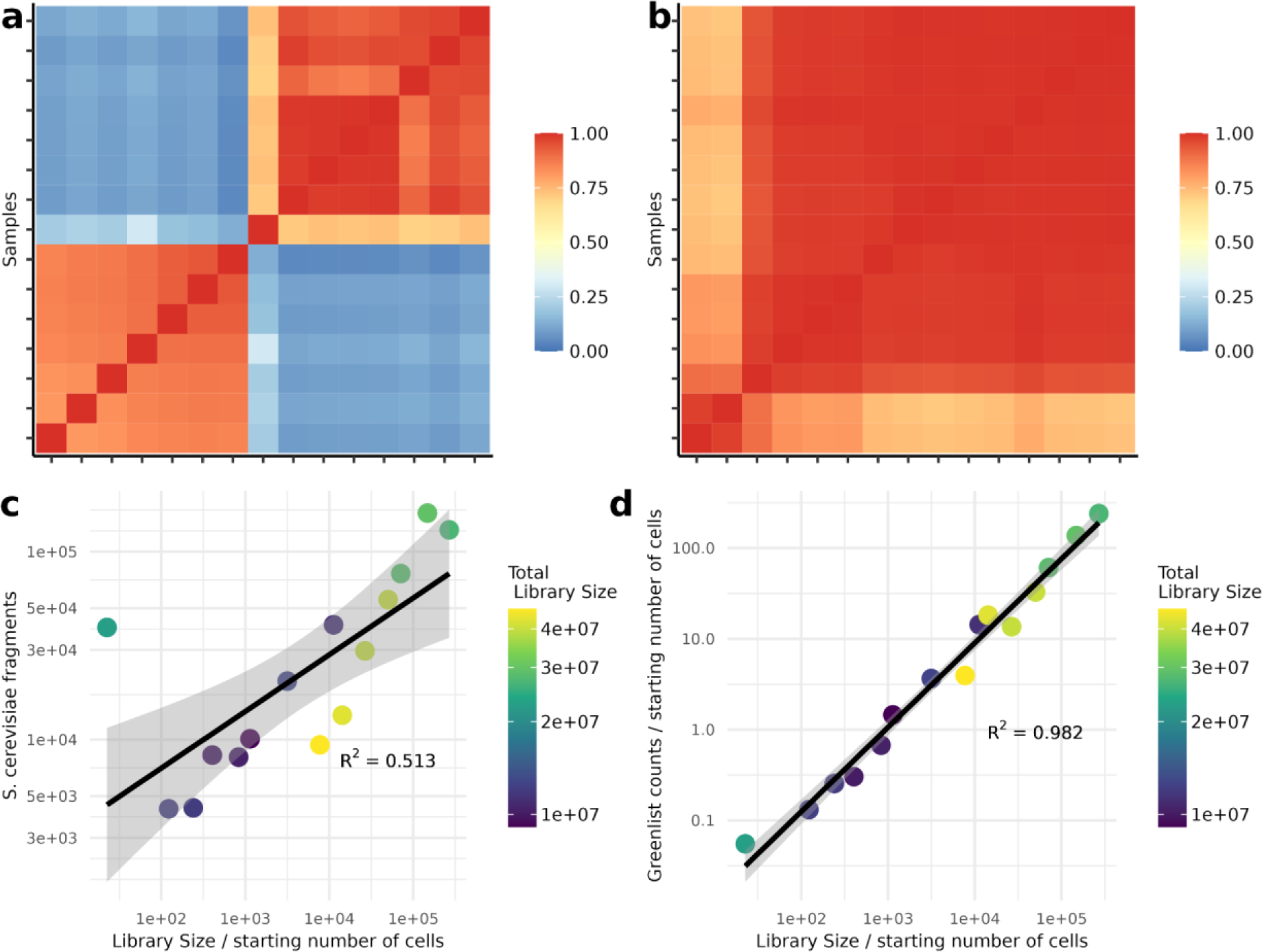
Greenlist normalization outperforms spike-in on dataset GSE104550 from the Henikoff lab. **a,** Heatmap of Pearson correlation between all samples, considering all 1kb genomic bins; **b,** Heatmap of Pearson correlation between all samples, considering only greenlist regions; **c,** Correlation between spike amplification (inferred as *S. cerevisiae* fragments reads / original fragments, where original spike in fragments remain constant for all samples) and *de facto* library amplification (inferred as total library size reads / starting number of cells); **d,** Correlation between greenlist amplification (inferred as total greenlist counts / starting number of cells) and *de facto* library amplification (as in **c** above). Linear regression and 95% confidence interval shown for graphs c and d, R^2^ calculated as fit to the linear regression.

The design of this experiment [26] proved very advantageous for testing normalization options, as the samples are expected to remain biologically consistent within each of the two antibodies tested, and the main sources of variation should be the starting number of cells (tested here at several values between 100 – 1,000,000 cells) and the technical variations expected from DNA extraction, library preparation, and sequencing. Thus, we can consider the ratio of library size to starting cell count as the *de facto* technical variation, as we expect around the same number of fragments generated per cell in each sample, and we can evaluate each normalization method as to how well it can account for this variation. We saw that spike-in normalization (either by *E. coli* carryover or added *S. cerevisiae* material) exhibited low correlations to our known *de facto* technical variations (R^2^=0.513) (Fig. 5c), failing to arrive at consistently normalized libraries. On the other hand, greenlist counts appropriately encapsulate this variation (R^2^=0.982) (Fig. 5d). Importantly, while the exact cell count is known in this experiment [26], as the variation was an intended design, it does not need to be known in advance for other experimental applications; thus, greenlist normalization can more accurately account for unknown variations in cell counts that may arise from experimental conditions.

Overall, these results demonstrate an intrinsic advantage of endogenous normalization factors such as our greenlist. As they are generated concurrently with the fragments of interest, they are directly affected by the same technical and experimental variations, such as different starting cell counts or variations in yield as discussed earlier.

### Greenlist normalization uncovers relevant variation from biased datasets

Lastly, we aimed to validate our greenlist’s applicability as a normalization factor in real experimental scenarios. For this, we specifically looked for CUT&RUN studies that featured extreme differences in library sizes, as they stand to benefit the most from a novel, robust normalization approach. We first selected study GSE221701 [27], which featured 48 samples and used 11 antibodies, with sample library sizes ranging from 11 to 134 million read pairs. Secondly, study GSE194217 [28], with 26 samples and 6 antibodies, with sample library sizes ranging from 3 to 64 million reads. Notably, both works evaluated the CUT&RUN results in only a qualitative manner, instead seeking quantitative validations from other techniques such as RNA-seq, ATAC-seq, and ChIP-seq.

To evaluate the impact of technical variation in the above datasets and evaluate the potential of proper normalization to mitigate it, we performed a principal component analysis (PCA) of the genomic data under different normalization scalings. Should the samples be improperly normalized, we expected the variations of large library size samples to contribute to the composition of PCs more strongly, thus biasing the graph and eclipsing variation patterns of interest (such as those related to biological sources). Applying standard normalization, based only on total library size, we saw that the PCAs of each dataset seemed to form clusters divided by antibodies (Fig. 6a-b), but there was a strong correlation between PC1 and library size (Fig. 6c-d). Thus, the variation observed was strongly impacted by technical variation, even after normalization, and attempting to interpret the data under these conditions could easily lead to inaccurate biological conclusions. Zou *et al*. [28], authors of dataset GSE194217, did perform spike-in normalization with added *S. cerevisiae* DNA; performing the PCA analysis with this normalized data, we saw that it did remove the correlation with library size (Extended Data 6a) – but only to replace it with a correlation to raw spike-in counts (Extended Data 6b), thus still arriving at biased results.

**Figure 6:**
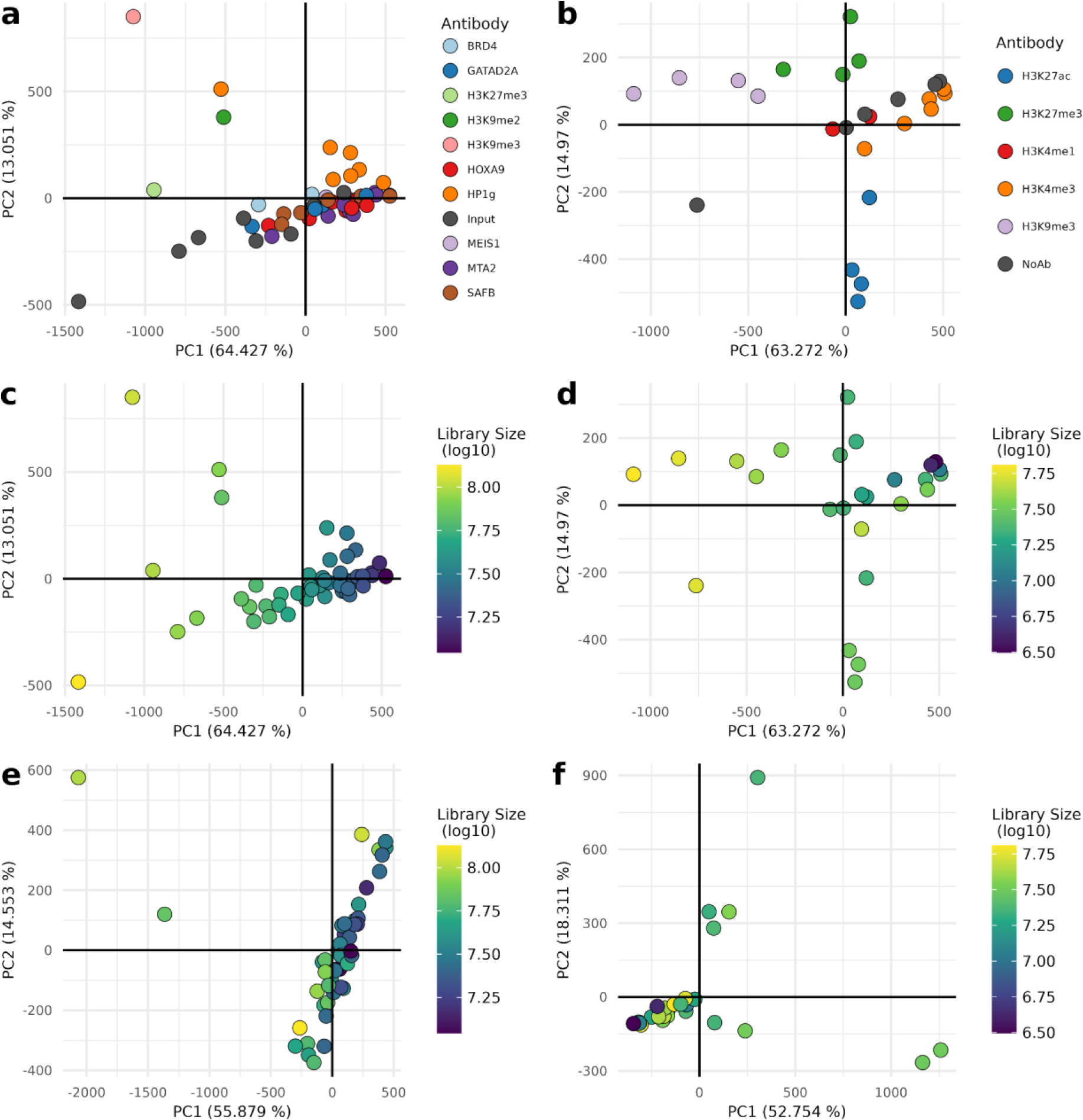
Greenlist normalization greatly minimizes technical variation, uncovering relevant biological data. Validation of greenlist normalization performance for two public datasets, GSE221701 (graphs **a**, **c**, **e**, Extended 6c, Extended 6e) and GSE194217 (graphs **b**, **d**, **f**, Extended 6a, Extended 6b, extended 6d, Extended 6f). **a–b,** First two components for PCA of each dataset, using default library size-based normalization; samples colored by antibodies used; **c–d,** First two components for PCA of each dataset, using default library size-based normalization, with samples colored by library sizes; **e– f,** First two components for PCA of each dataset, calculated after performing greenlist normalization; samples colored by library sizes.

On the other hand, performing the analysis on greenlist-normalized counts greatly minimized the bias of library size (Fig. 6e-f), without creating a bias to the greenlist itself (Extended Data 6c-d). As such, the analysis can now be appropriately interpreted in a biological context as a typical PCA would be (Extended Data 6e-f). For the work of Zou *et al.* in particular, this new analysis now highlights two histone marks, H3K27ac and H3K27me3 (Extended Data 6f), which is in line with some of the major findings of their work [28].

## Discussion

High-throughput genomic techniques such as CUT&RUN can be especially sensitive to poor normalization, as the vastly different signal profiles of different targets and experimental designs can interfere with typical statistical assumptions. Consequently, this poor normalization risks jeopardizing the technique’s quantitative power. And as new experimental protocols are developed, so too do specialized *in silico* tools become necessary for their proper analysis. In that sense, we here introduce a new normalization technique, highly specialized for either CUT&RUN or CUT&Tag, through a novel application of information theory approaches to the underlying concept of the ENCODE ChIP-seq blacklist [21]. Our approach leverages the extensive collection of datasets published thus far, and through rigorous testing and validation we observed our greenlist’s applicability across a variety of experimental scenarios. Furthermore, we showed that greenlist normalization outperforms the current standards. This offers a robust analysis option, which should apply to any CUT&RUN or CUT&Tag experiment, with no added experimental step, and incurring no additional costs. Even for organisms featuring few published datasets, we showed that this entropy-based greenlist construction should remain consistent and still offer a robust normalization. We expect that this methodology will enhance the quantitative potential of CUT&RUN and CUT&Tag in the scientific literature, fostering more comprehensive and reliable analyses.

## Supporting information

Supplementary Materias S1

Supplementary Materias S2

Supplementary Materias S3

Supplementary Materias S4

## Extended Data

**Extended Data 4:**
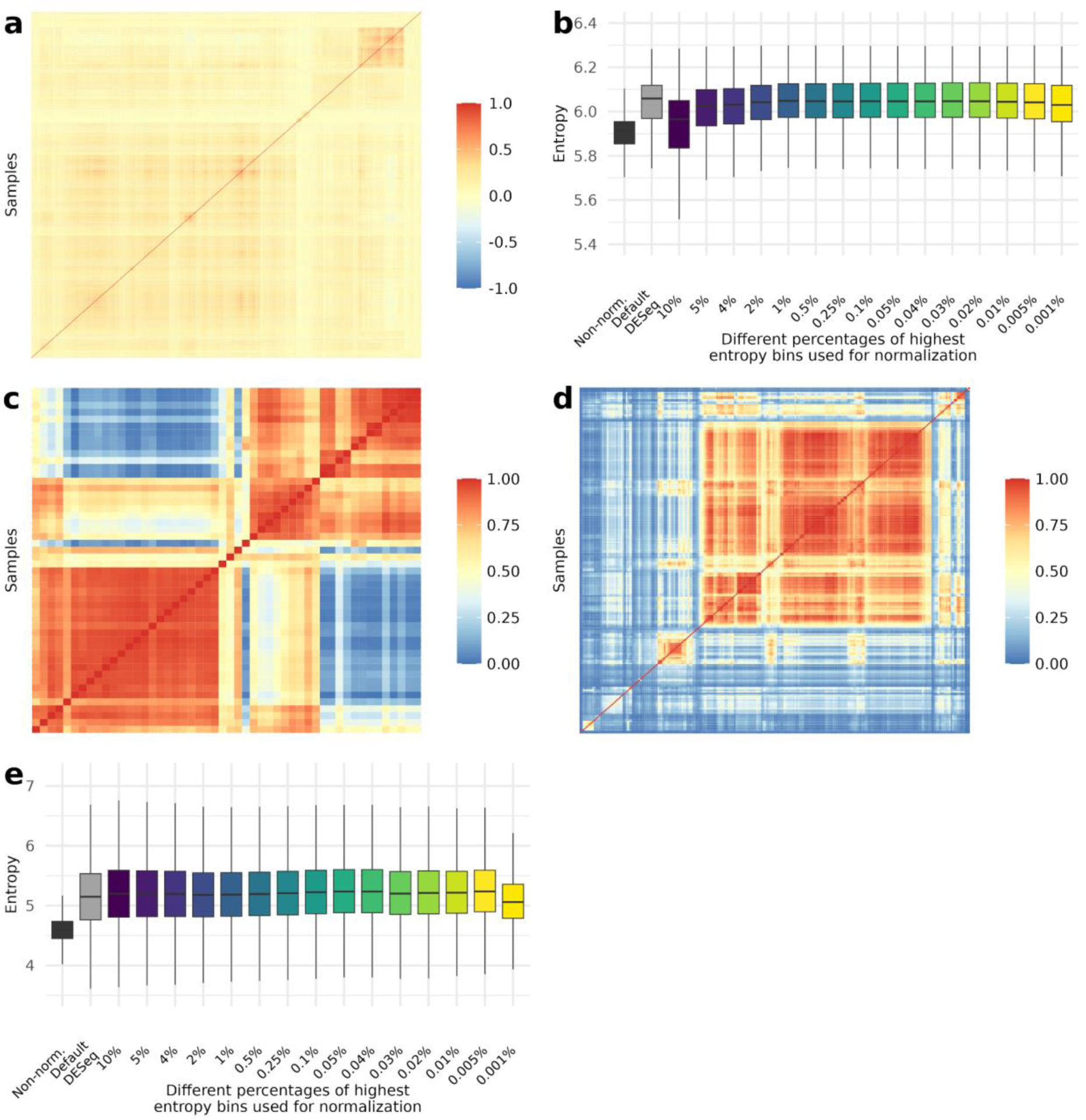
Other applications of our pipeline show similar performances. **a,** Heatmap of Pearson correlation of 611 mouse CUT&RUN samples, considering signal from all 1kb genomic bins; **b,** Entropy distributions for our mouse CUT&RUN test set after normalization using different percentages of highest entropy bins, as indicated in the x-axis; in the y-axis, boxplot represents median entropy, first and third quartile, whiskers extend to 5% and 95% quantiles; **c,** Heatmap of Pearson correlation of the 50 samples from dataset GSE151326, considering signal from all 1kb genomic bins; **d,** Heatmap of Pearson correlation of 217 human CUT&Tag samples, considering signal from all 1kb genomic bins; **e,** Entropy distributions for our human CUT&Tag test set after normalization using different percentages of highest entropy bins, as indicated in the x-axis; in the y-axis, boxplot represents median entropy, first and third quartiles, whiskers extend to 5% and 95% quantiles.

**Extended Data 6:**
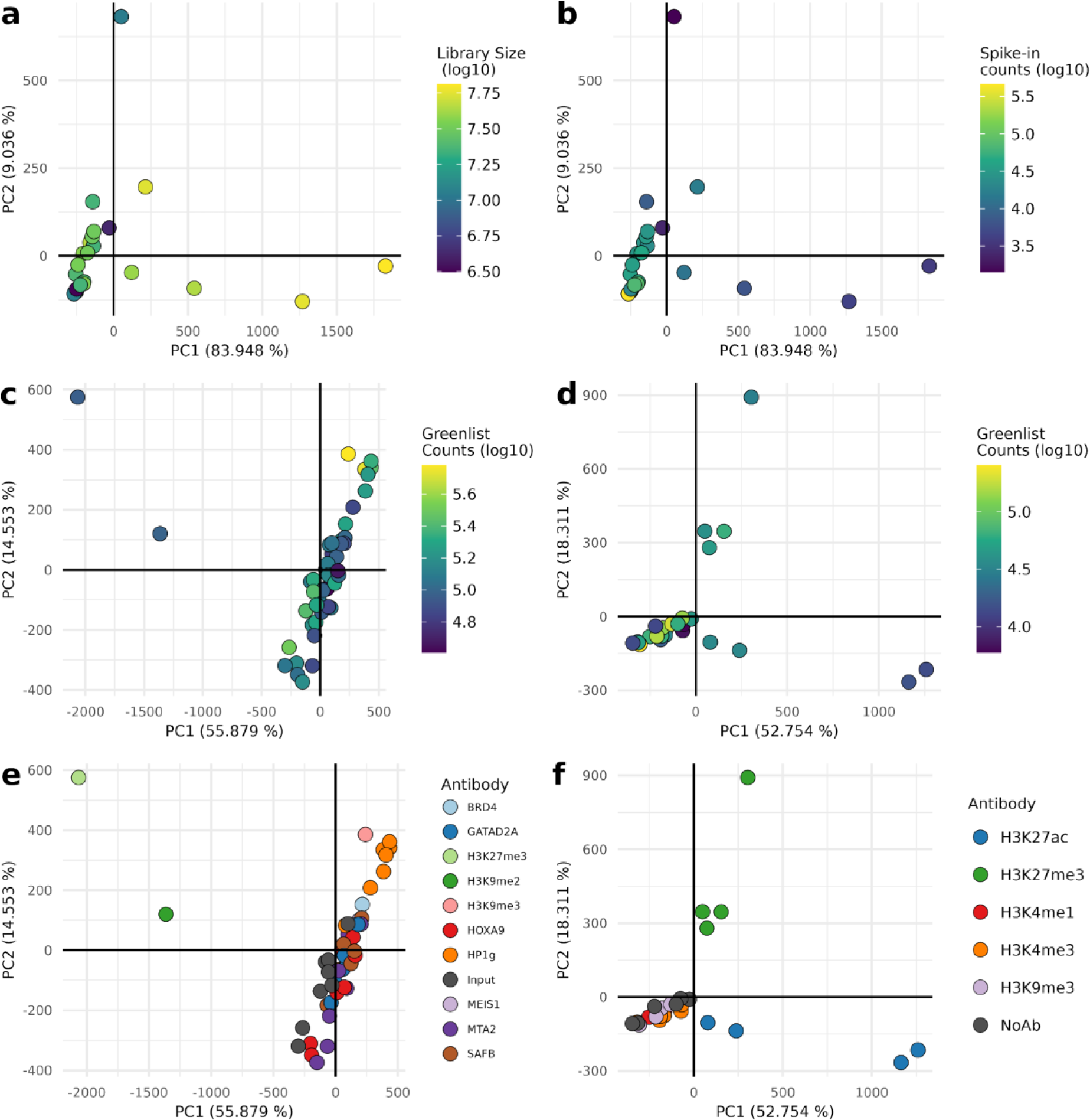
Greenlist normalization greatly minimizes technical variation, uncovering relevant biological data. Validation of greenlist normalization performance for two public datasets, GSE221701 (graphs **a**, **c**, **e**, Extended 6c, Extended 6e) and GSE194217 (graphs **b**, **d**, **f**, Extended 6a, Extended 6b, extended 6d, Extended 6f). **a–b,** First two components for PCA of dataset GSE194217, using spike-in normalization; samples colored by library sizes (**a**) or spike-in counts (**b**); **c–d,** First two components for PCA of each dataset, calculated after performing greenlist normalization, with samples colored by total counts in greenlist regions; **e–f,** First two components for PCA of each dataset, calculated after performing greenlist normalization; samples colored by antibodies used.

## Online Methods

### Sample selection and processing

Samples were selected from publicly available studies submitted to the Sequence Read Archive repository (SRA, https://www.ncbi.nlm.nih.gov/sra), and manually curated to ensure the consistency and accuracy of the metadata. Experimental meta-factors considered included cell line, cell type/characteristics, antibody brand and product, antibody target (eg. anti-mouse IgG versus anti-human IgG), antibody host, CUT&RUN protocol/kit used, authors and authors’ primary affiliations. Only negative control samples from studies with ≥ 6 samples total were selected. Samples with too few aligned reads were discarded (≤ 1.5M for human CUT&RUN, ≤ 1M for mouse CUT&RUN, ≤ 500k for human CUT&Tag), based on recommendations of established protocols [6, 29–31]. In total, 463 samples were used for human CUT&RUN, 611 for mouse CUT&RUN and 217 for human CUT&Tag; a full list of datasets used is available in Supplementary Material S4.

Samples were downloaded with SRA-toolkit (SRA Toolkit Development Team, v3.0.2 https://trace.ncbi.nlm.nih.gov/Traces/sra/sra.cgi?view=software), cleaned with fastp (v0.20.0) [32] with parameters “--detect_adapter_for_pe -W 1 -3 -5”, and aligned with bowtie2 (v2.3.5.1) [33] with parameters “-X 1000 --no-mixed --dovetail --no-unal --very-sensitive-local -N 1”. Genome builds used were hg38/GRCh38 with gene annotation GENCODE Release 40 [34] for human, mm39/GRCm39 with gene annotation GENCODE Release M33 [34] for mouse, R64-1-1 (Saccharomyces Genome Database, https://www.yeastgenome.org/) for yeast, and DH5alpha strain for *E. coli* (GenBank entry CP026085.1). Files were processed as needed with samtools (v1.10) [35] and bedtools (v2.26.0) [36].

For the validation analyses of specific experiments (GSE104550, GSE221701, and GSE194217), we followed their original published methodology for sample processing [26–28].

### Construction of the greenlist

Binning was performed with 1kb windows while blacklisting only the “Low-Mappability” regions of the ENCODE blacklist [21]. Quantifications were done with deeptools (v3.5.1) [37] multiBamSummary, either in bins mode for bins or bed-file mode for quantifying finished greenlists. Quantified bins were normalized by quantile normalization with the broman (v0.80) R package [38], and Shannon entropy was calculated with the entropy (v1.3.1) R package [39], using the maximum-likelihood approximation. Briefly, this metric quantifies the expected value of the information content of a variable; in this case, for each bin X, we consider the proportion of counts per sample over the total counts of X as the probability to calculate X’s information content, so that bins with high-count outliers have low information entropies, and bins with a more homogenous distribution have high entropies. For calculating normalization factors, we tested both a manual approach (simply dividing (library size)/sum(greenlist counts)) or using DESeq2 (v1.32.0) [40] to calculate size factors; both approaches showed similar results, so DESeq2 was used for convenience.

After testing, final constructions of the greenlists were done by selecting bins in the top 0.1% of highest entropies, filtering out bins < 5kb away from known genes, and extending them if the bins in their proximity were in the top 1% of highest entropies. Blacklist constructions were done in a similar fashion but selecting bins with median normalized counts in the top 0.1% and 1% of highest signal instead.

In the interest of reproducibility, the R scripts used for greenlist construction and validation are provided at https://github.com/fndemello/CUT-RUN_greenlist.

### Validations and statistics

For comparisons between lists, correlation was calculated as Spearman’s rank correlation coefficient, ranking bins by entropy (for greenlists) or highest signal (for blacklists). Overlap comparisons were calculated by finding intersects with bedtools [36] and calculating F-scores for each pair of lists.

The Monte-Carlo test to validate our analysis of normalization efficiency of different thresholds (Fig. 2c, 3b) was done with base R, the doParallel (v1.0.16) package for parallelization and doRNG (v1.8.6) package to ensure random seed consistency across worker threads. Tests were done by selecting 0.1% of bins (2789) with the sample function and testing their effect on entropy of our test set (10% of lowest entropy bins, as previously), with 100,000 trials. Efficiency was also assessed after filtering out sampled regions that were close/overlapping gene bodies.

For analyzing the association of meta-factors to the overall variation of the datasets, PCA was performed, and a linear regression model was built for each of the first 5 PCs. Significance was tested with ANCOVA F-test, with a threshold of p ≤ 0.01 for significance with the R package car (v3.0-12) [41]. It is worth noting that this statistical design is by nature unbalanced, as we cannot ensure that all categorical variables are equally represented without greatly downsampling our dataset. As such, some groups are incomplete (i.e. not every combination of variable values is present) and some values have a small representation (e.g. antibodies that were only used once across the whole dataset), slightly limiting the estimations; nevertheless, ANCOVA should be resilient enough to this imbalance.

Additional statistical analyses were done with R (v4.1.0) [42], and plotting was done with ggplot2 (v3.4.2) [43].

## Notes

https://github.com/fndemello/CUT-RUN_greenlist

### Competing Interest Statement

The authors have declared no competing interest.

## References

1. Johnson, D.S., et al., Genome-wide mapping of in vivo protein-DNA interactions. Science, 2007. 316(5830): p. 1497–502.

2. Park, P.J., ChIP-seq: advantages and challenges of a maturing technology. Nat Rev Genet, 2009. 10(10): p. 669–80.

3. Furey, T.S., ChIP-seq and beyond: new and improved methodologies to detect and characterize protein-DNA interactions. Nat Rev Genet, 2012. 13(12): p. 840–52.

4. Skene, P.J. and S. Henikoff, An efficient targeted nuclease strategy for high-resolution mapping of DNA binding sites. Elife, 2017. 6: e21856.

5. Schmid, M., T. Durussel, and U.K. Laemmli, ChIC and ChEC; genomic mapping of chromatin proteins. Mol Cell, 2004. 16(1): p. 147–57.

6. Meers, M.P., et al., Improved CUT&RUN chromatin profiling tools. Elife, 2019. 8: e46314.

7. Meers, M.P., D. Tenenbaum, and S. Henikoff, Peak calling by Sparse Enrichment Analysis for CUT&RUN chromatin profiling. Epigenetics Chromatin, 2019. 12(1): p. 42.

8. Salma, M., et al., High-throughput methods for the analysis of transcription factors and chromatin modifications: Low input, single cell and spatial genomic technologies. Blood Cells Mol Dis, 2023. 101: p. 102745.

9. Klein, D.C. and S.J. Hainer, Genomic methods in profiling DNA accessibility and factor localization. Chromosome Res, 2020. 28(1): p. 69–85.

10. Leo, L. and N. Colonna Romano, Emerging Single-Cell Technological Approaches to Investigate Chromatin Dynamics and Centromere Regulation in Human Health and Disease. Int J Mol Sci, 2021. 22(16): p. 8809.

11. Kaya-Okur, H.S., et al., CUT&Tag for efficient epigenomic profiling of small samples and single cells. Nat Commun, 2019. 10(1): p. 1930.

12. Meers, M.P., D.H. Janssens, and S. Henikoff, Pioneer Factor-Nucleosome Binding Events during Differentiation Are Motif Encoded. Mol Cell, 2019. 75(3): p. 562–575 e5.

13. Yu, F., V.G. Sankaran, and G.C. Yuan, CUT&RUNTools 2.0: a pipeline for single-cell and bulk-level CUT&RUN and CUT&Tag data analysis. Bioinformatics, 2021. 38(1): p. 252–254.

14. Boyd, J., et al., ssvQC: an integrated CUT&RUN quality control workflow for histone modifications and transcription factors. BMC Res Notes, 2021. 14(1): p. 366.

15. Orlando, D.A., et al., Quantitative ChIP-Seq normalization reveals global modulation of the epigenome. Cell Rep, 2014. 9(3): p. 1163–70.

16. Ghosh, D. and Z.S. Qin, Statistical Issues in the Analysis of ChIP-Seq and RNA-Seq Data. Genes (Basel), 2010. 1(2): p. 317–34.

17. Dickson, B.M., et al., A physical basis for quantitative ChIP-sequencing. J Biol Chem, 2020. 295(47): p. 15826–15837.

18. Chen, K., et al., The Overlooked Fact: Fundamental Need for Spike-In Control for Virtually All Genome-Wide Analyses. Mol Cell Biol, 2015. 36(5): p. 662–7.

19. Grzybowski, A.T., Z. Chen, and A.J. Ruthenburg, Calibrating ChIP-Seq with Nucleosomal Internal Standards to Measure Histone Modification Density Genome Wide. Mol Cell, 2015. 58(5): p. 886–99.

20. Bonhoure, N., et al., Quantifying ChIP-seq data: a spiking method providing an internal reference for sample-to-sample normalization. Genome Res, 2014. 24(7): p. 1157–68.

21. Amemiya, H.M., A. Kundaje, and A.P. Boyle, The ENCODE Blacklist: Identification of Problematic Regions of the Genome. Sci Rep, 2019. 9(1): p. 9354.

22. Carroll, T.S., et al., Impact of artifact removal on ChIP quality metrics in ChIP-seq and ChIP-exo data. Front Genet, 2014. 5: p. 75.

23. Wimberley, C.E. and S. Heber, PeakPass: Automating ChIP-Seq Blacklist Creation. J Comput Biol, 2020. 27(2): p. 259–268.

24. Stark, R. and G.D. Brown. DiffBind: differential binding analysis of ChIP-Seq peak data. 2011; Available from: http://bioconductor.org/packages/release/bioc/html/DiffBind.html.

25. Nordin, A., et al. The CUT&RUN suspect list of problematic regions of the genome. Genome Biol, 2023. 24: p. 185.

26. Skene, P.J., J.G. Henikoff, and S. Henikoff, Targeted in situ genome-wide profiling with high efficiency for low cell numbers. Nat Protoc, 2018. 13(5): p. 1006–1019.

27. Agrawal-Singh, S., et al., HOXA9 forms a repressive complex with nuclear matrix-associated protein SAFB to maintain acute myeloid leukemia. Blood, 2023. 141(14): p. 1737–1754.

28. Zou, H., et al., A neurodevelopmental epigenetic programme mediated by SMARCD3-DAB1-Reelin signalling is hijacked to promote medulloblastoma metastasis. Nat Cell Biol, 2023. 25(3): p. 493–507.

29. Kong, N.R., et al., A modified CUT&RUN protocol and analysis pipeline to identify transcription factor binding sites in human cell lines. STAR Protoc, 2021. 2(3): p. 100750.

30. EpiCypher. Available from: https://www.epicypher.com/content/documents/protocols/cutana-cut&run-protocol-2.1.pdf.

31. Cell_Signaling_Technology. Available from: https://www.cellsignal.com/learn-and-support/protocols/cut-and-run-protocol.

32. Chen, S., et al., fastp: an ultra-fast all-in-one FASTQ preprocessor. Bioinformatics, 2018. 34(17): p. i884–i890.

33. Langmead, B. and S.L. Salzberg, Fast gapped-read alignment with Bowtie 2. Nat Methods, 2012. 9(4): p. 357–9.

34. Frankish, A., et al., Gencode 2021. Nucleic Acids Res, 2021. 49(D1): p. D916–D923.

35. Danecek, P., et al., Twelve years of SAMtools and BCFtools. Gigascience, 2021. 10(2): p. giab008.

36. Quinlan, A.R. and I.M. Hall, BEDTools: a flexible suite of utilities for comparing genomic features. Bioinformatics, 2010. 26(6): p. 841–2.

37. Ramirez, F., et al., deepTools2: a next generation web server for deep-sequencing data analysis. Nucleic Acids Res, 2016. 44(W1): p. W160–5.

38. Broman, K. and J. Tian. kbroman/broman: Version 0.80 (0.80) Zenodo. 2022; Available from: 10.5281/zenodo.6811647.

39. Hausser, J. and K. Strimmer, Entropy Inference and the James-Stein Estimator, with Application to Nonlinear Gene Association Networks. Journal of Machine Learning Research, 2009. 10: p. 1469–1484.

40. Love, M.I., W. Huber, and S. Anders, Moderated estimation of fold change and dispersion for RNA-seq data with DESeq2. Genome Biol, 2014. 15(12): p. 550.

41. Fox, J. and S. Weisberg. An R Companion to Applied Regression, Third edition. 2019; Available from: https://socialsciences.mcmaster.ca/jfox/Books/Companion/.

42. R_Core_Team. R: A language and environment for statistical computing. 2021; Available from: https://www.R-project.org/.

43. Wickham, H., ggplot2: Elegant Graphics for Data Analysis. 2016. 2nd ed., Springer, ISBN 978-3319242750.

