## Supplementary Materias S3 for "The CUT&RUN Greenlist: genomic regions of consistent noise are effective normalizing factors for quantitative epigenome mapping"

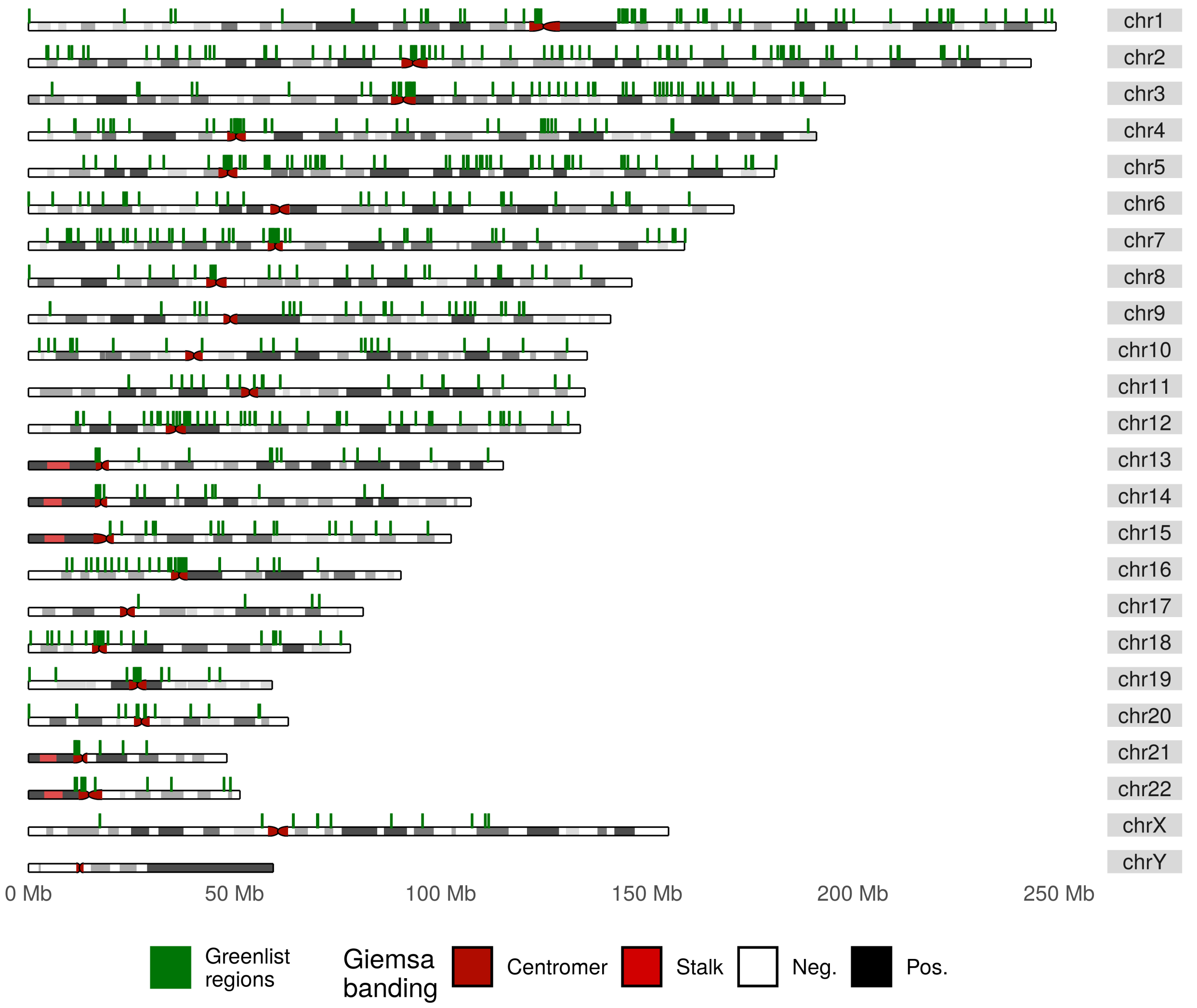


**Figure 1:** Distribution of CUT&RUN Greenlist regions (green) across the human genome. Typical karyotype Giemsa staining represented for reference.

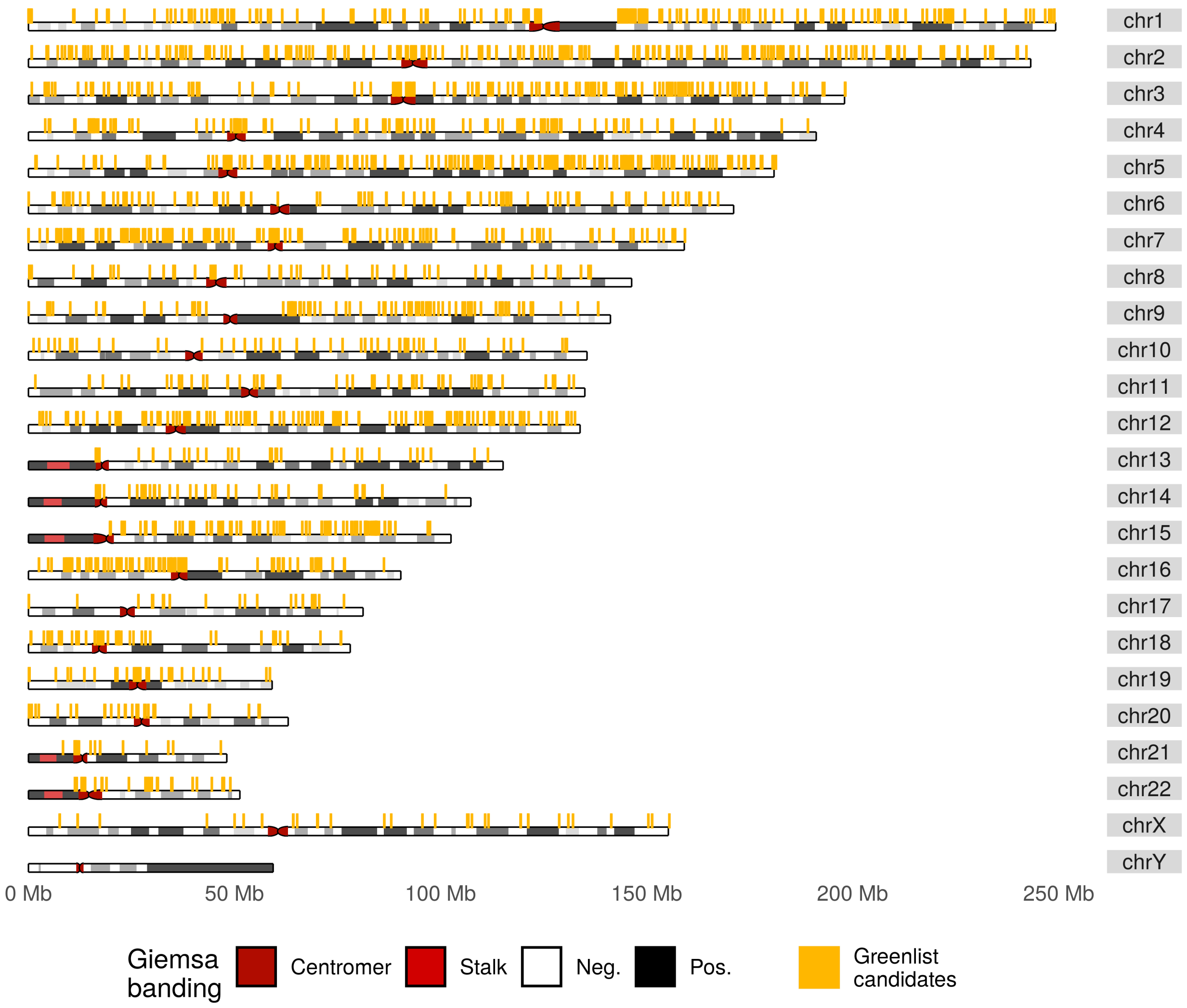


**Figure 2:** Distribution of candidate CUT&RUN Greenlist regions (yellow) across the human genome, including those later excluded for overlapping genic regions. Typical karyotype Giemsa staining represented for reference.

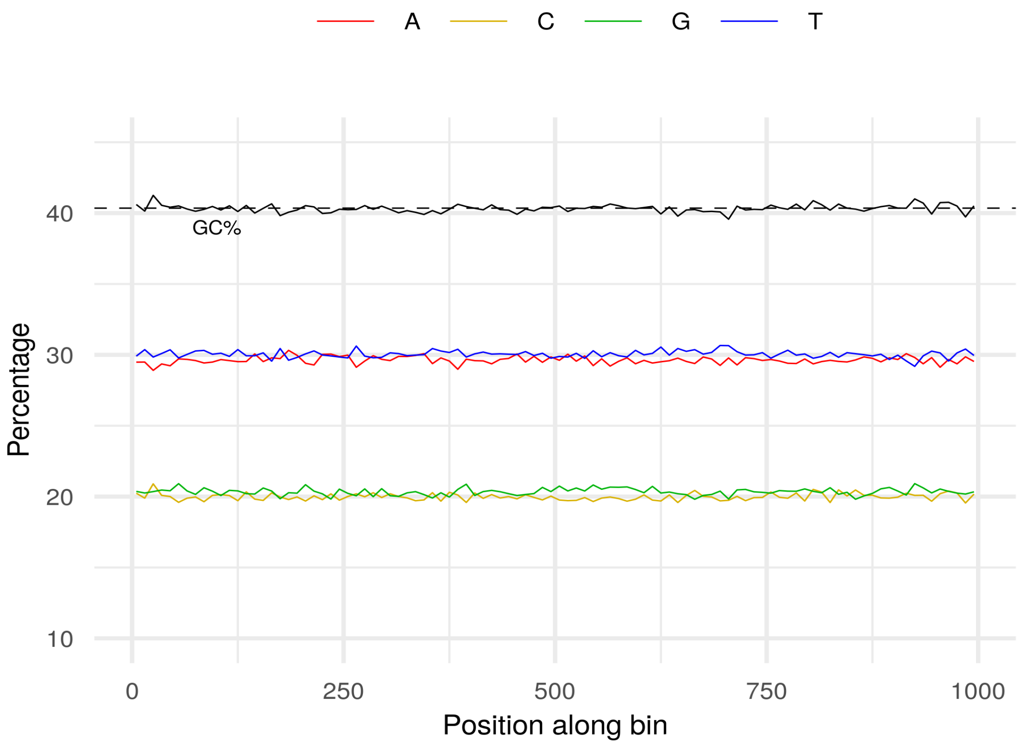


**Figure 3:** Frequency distribution of nucleotides along bins selected for the human CUT&RUN Greenlist. GC% represented in black, with overall average GC% shown as a dashed line.

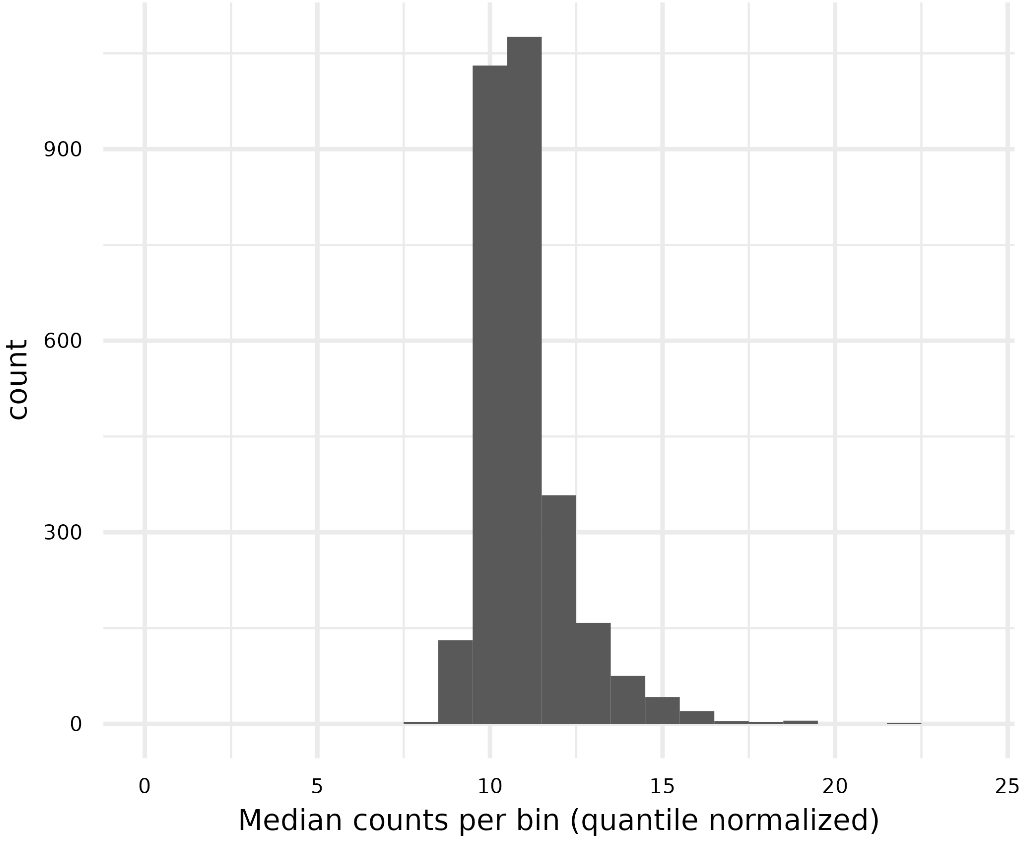


**Figure 4:** Median normalized counts per bin for the human CUT&RUN Greenlist, 0-99% quantile displayed. High-count outliers (40 regions with 1000-5500 counts each, which correspond to the overlap between the CUT&RUN Greenlist and Blacklist) excluded to avoid distorting x-axis scaling.

**Table 1:** Distribution of human CUT&RUN Greenlist regions over genomic features of interest, compared to whole genome distributions.

| Feature | Greenlist candidate bins | Selected Greenlist bins | All genomic bins |
|---|---|---|---|
| Bins overlapping gene bodies & gene neighborhoods | 36.148 % | 0 % | 31.195 % |
| Bins overlapping centromer regions | 6.850 % | 18.949 % | 2.032 % |
| Bins overlapping euchromatin* | 42.557 % | 41.557 % | 52.270 % |
| Bins overlapping heterochromatin* | 48.389 % | 35.460 % | 43.955 % |

*Defined as positive (euchromatin) or negative (heterochromatin) consensus Giemsa band staining (NCBI GRCh38.p12 cytogenetic annotation).

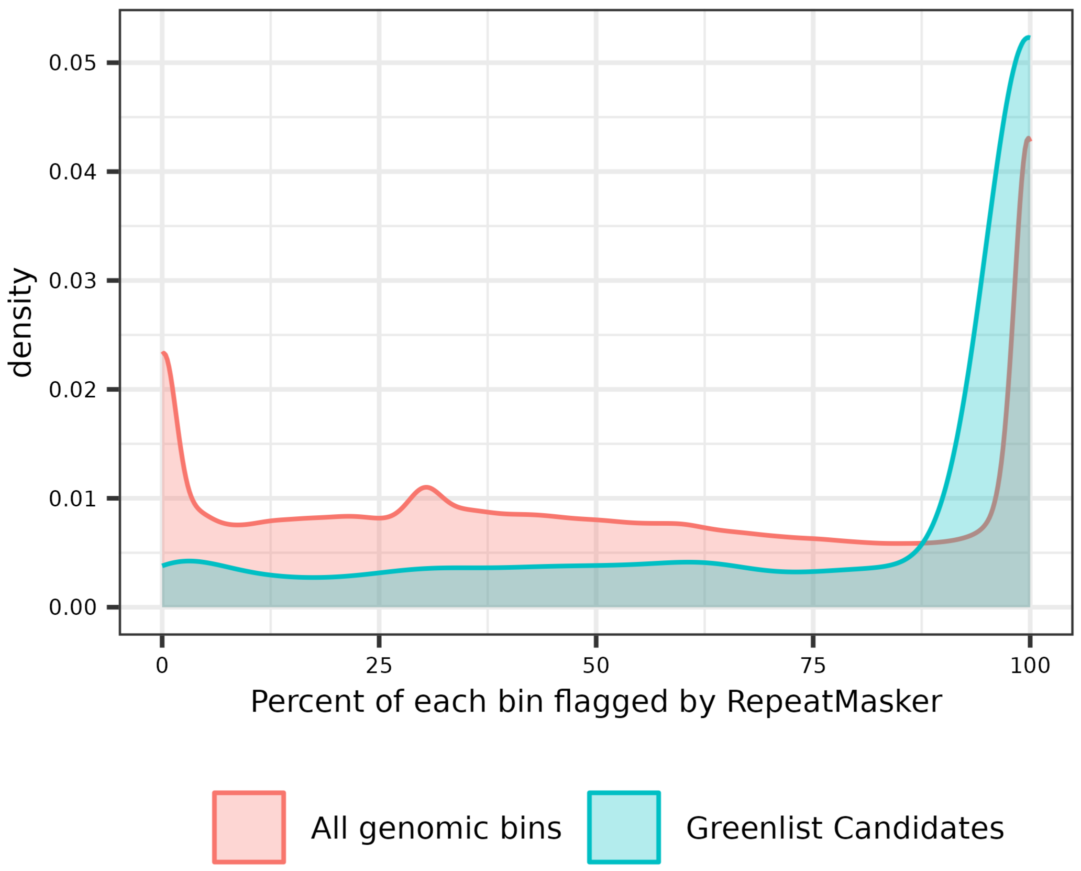


**Figure 5:** Density distribution of percentage of repetitve elements (identified by RepeatMasker) per 1kb bin, either for all genomic bins (red) or high entropy bins used for greenlist creation (blue).

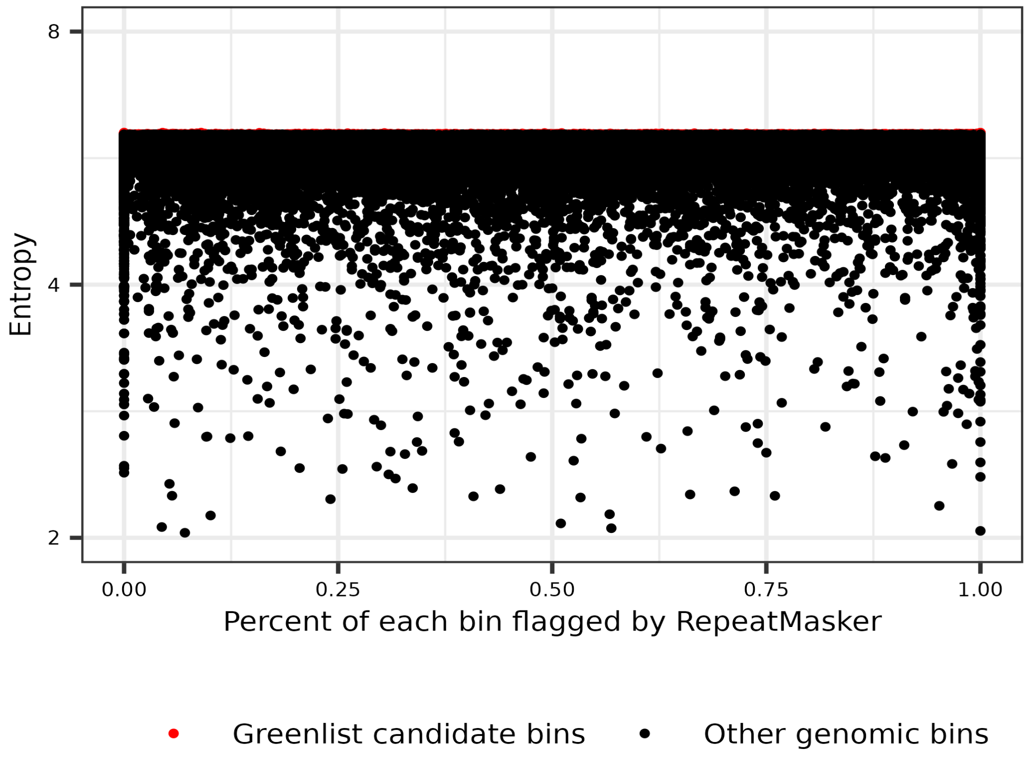


**Figure 6:** Distribution of 1kb human genomic bins by percentage of repetitve elements identified by RepeatMasker (x-axis) and calculated Shannon entropy for the human CUT&RUN samples (y-axis). Bins identified as either Greenlist candidates (red) or non-cadidates (black).
